# Effects of substrate on shrimp growth, water quality and bacterial community in the biofloc system nursing *Penaeus vannamei* postlarvae

**DOI:** 10.1101/2022.02.08.479639

**Authors:** Hai-Hong Huang, Chao-Yun Li, Yan-Ju Lei, Wei-Qi Kuang, Bo-Lan Zhou, Wan-Sheng Zou, Pin-Hong Yang

## Abstract

This study aimed to investigate the effects of substrate on water quality, shrimp growth and bacterial community in the biofloc system with a salinity of 5‰. Two treatments, biofloc system with (sB) or without (nB) addition of elastic solid packing filler (nylon) as substrate, were set up. *Penaeus vannamei* postlarvae (PL, ~ stage 15) were stocked at a density of 4000 PL m^−3^ in each treatment with triplicates for a 28-days culture experiment, taking glucose as carbon source (C:N 15:1). Results showed that the survival rate (96.3±3.6%), FCR (0.76±0.06) and productivity (1.54±0.12 kg m^−3^) in sB treatment were significantly better than those in nB treatment (81.0±7.1%, 0.98±0.08 and 1.14±0.09 kg m^−3^, *P*<0.05). All water parameters were in the recommended ranges. Substrate showed significant effect on TAN, TSS, turbidity, biofloc volume, pH and carbonate alkalinity (*P* < 0.05). Actinobacteria (4.0-22.7%), Bacteroidetes (10.4-33.5%), Firmicutes (0.2-11.2%), Planctomycetes (4.0-14.9%) and Proteobacteria (29.4-59.0%) were the most dominant phyla for both treatments. However, the bacterial community in sB treatment showed to be significantly different from that in nB treatment (Jaccard distance 0.94±0.01, *P*=0.001). Substrate showed significant effects on Shannon, Heip, Pielou and Simpson index, as well as relative abundances of Actinobacteria, Bacteroidetes and Planctomycetes (*P* < 0.05). The results suggested that addition of substrate affected the shrimp growth, water quality and bacterial community in the biofloc system nursing *P. vannamei* PL with a 5‰ salinity.

## 1 INTRODUCTION

Crustacean is the most valuable group in fish trade in the world, accounting for 23.0% in terms of value with an 8.3% share in culture production, compared to the 65.4% Vs. 79.8% of fish, and the 11.6% Vs. 11.9% of mollusks and other aquatic invertebrates (FAO, 2018). Among the cultured crustaceans, *Penaeus vannamei* (Boone, 1931) is the most important species whose production accounted for 53.0% of the total crustaceans production (FAO, 2018).

Prenursery of postlarvae (PL) is a very important step during culture of *P. vannamei* in many places (Mishra et al., 2008; Rezende et al., 2018). In the conventional prenursery system, PL are intensively stocked at a high density (up to 60 PL L^−1^) with small-scale water volume and carefully nursed for 15-60 days (Schveitzer et al., 2017), to save water, land and energy resources, and to deliver large individuals that are resistant to environmental conditions to increase the survival and growth performance during the grow-out phase (Mishra et al., 2008; Rezende et al., 2018). However, in this high-stocking-density system, serious cannibalism behavior, competition for space and degradation of water quality would affect growth and survival of PL (Esparza-Leal et al., 2015). The use of artificial substrate is an effective approach to supply additional surface and refugee for shrimp during prenursery (Tierney et al., 2020). Additionally, biofloc technology (BFT) has also been tried to be used to prenursery of *P. vannamei* PL (Khanjani et al., 2017; Schveitzer et al., 2017), to control the problems usually appeared in the traditional prenursery system, such as biosecurity and toxic ammonia and nitrite (Samocha, 2010), due to the advantages of this technology on nitrogen assimilation *in situ* and pathogen control under minimal or zero water exchange conditions (Avnimelech, 2015; Huang, 2020). Furthermore, several studies have used substrates in biofloc systems to reinforce the controlling of water quality and the growth performance of PL by supplying a natural food through the periphyton growing on the substrate during the prenursery stage (Peiro-Alcantar et al., 2019; Rezende et al., 2018; Tierney et al., 2020).

*P. vannamei* is an euryhaline species, and can be cultured at low salinity conditions less than 1.0‰, which is a trend that will continue to grow globally (Roy et al., 2010). Osmoregulation capability of *P. vannamei* develops gradually in the postlarvae stages and can be acclimated to salinities as low as 0.5‰ by stage 12 (Van Wyk et al., 1999), indicating that PL could be nursed under low salinity conditions. As well, biofloc technology has been successfully used for prenursery of *P. vannamei* PL under salinity conditions of 8-16‰ (Esparza-Leal et al., 2016; Luo et al., 2019). However, application of substrate in prenursery of *P. vannamei* PL in the biofloc system under a low-salinity condition has been not documented yet.

In this study, it is aimed to estimate the effects of substrate on shrimp growth performance and water quality in the biofloc system nursing *P. vannamei* postlarvae (PL) under a salinity condition as low as about 5‰. Furthermore, the effects of substrate on the bacterial community diversity and composition in this system were also investigated.

## 2 MATERIAL AND METHODS

### 2.1 Ethic statement

The culture experiments were carried out at a local farm of Bifuteng eco-agriculture development Co., Ltd. (BEAD Co., Ltd., Changde, China, Lat. 28°53’57.88” N, Long. 111°38’3.08” E), with indoor tanks, under principles in good laboratory animal care, according to the national standard of China (GB/T 35892-2018), ‘Laboratory animal-Guideline for ethical review of animal welfare’. The manuscript does not require ethical approval.

### 2.2 Experimental design

There two treatments with three replicates for each were set up in the present study, biofloc system with (sB) and without (nB) addition of substrate. In the sB treatment, 52.4% of the tank internal surface was equally coated with elastic solid packing filler (nylon, with a diameter of 15 cm).

Before the beginning of the experiment, tanks (width X length X depth = 2 × 2.5 × 1.3 m, 6.5 m^3^ in volume) connected with a Roots blower air pump (CRS-100, Changsheng mechanical and electrical equipment Co., Ltd., Zhangqiu, Shandong, China) were filled 5.0 m^3^ (1.0 m in depth) culture water with a salinity of about 5.0‰ per tank. The water for culture experiment was prepared according to the method of Ray and Lotz (2017), with some modifications. Briefly, artificial sea salt powder (Qianglong corporation, Tianjin, China), and food-grade chemical reagents of KCl, MgCl_2_, CaCl_2_ and NaHCO_3_ were added to tap water, to manage an initial salinity of 5.02±0.08‰, with K^+^, Mg^2+^ and Ca^2+^ concentrations of 301.3±5.9, 925.2±12.1 and 306.7±6.9 mg L^−1^, respectively, and a pH value of about 8.0. After that, sterilization with 10.0 mg L^−1^ of chlorinedioxide followed by neutralization with 1.0 mg L^−1^ of ascorbic acid was executed according to the processes of previous studies (Gaona et al., 2017; Lara et al., 2017).

At the beginning of the experiment, *P. vannamei* PL was randomly assigned to each tank at a density of 4000 PL per m^3^. Those PL (~PL15, 2.5±0.4 mg) were kindly supplied from BEAD Co., Ltd. and had been treated with a desalinating and acclimation procedure to adapt the experimental conditions before being assigned (Luo et al., 2019; Van Wyk et al., 1999). Next, during the 28-days experimental period, PL were fed with a commercial formulated shrimp diet (crude protein 40.0%, crude lipid 5.0%, crude fibber 5.0%, crude ash 15.0%, moisture 12.0%, Alpha corporation, Jiangmen, Guangdong, China), with a frequency of four times (6:00, 12:00, 18:00, 24:00) a day, basing on the feeding table and the estimated total biomass according to methods of Van Wyk et al. (1999). Besides, glucose (food grade, carbohydrate content 90.0%, Fufeng biotechnology Co., Ltd., Hohhot, Inner Mongolia Autonomous, China) was added as exogenous carbon source to the experimental tanks, according to an inputted carbon to nitrogen ratio (C:N) of 15:1 (Huang et al., 2021). The inputted C:N was the C:N contained in the inputted materials (feed and carbon source), and determined basing on the carbon contents contained in carbon source and feed (Kumar et al., 2017, Tinh et al. 2021) and the conception that 25% of the feed nitrogen is theoretically converted as shrimp biomass (Piedrahita, 2003). Throughout the whole experimental period, no water exchange was operated, but evaporating loss was complemented with dechlorinated tap water per week.

During 14-28 d of the experiment, partial biofloc was removed by using a side-stream settling chamber equipped a submersible pump (JGP-2500L, SENSEN Group Co., Ltd., Zhoushan, China) which ran for 6 h every day at a rate of 2.25 m^3^ per h to maintain a biofloc volume < 15 mL L^−1^.

### 2.3 Determination of water parameters

Water temperature, dissolved oxygen and pH were detected each day by using electric analyzers (YSI-ProPlus, Yellow Springs Instruments Inc., OH, USA; pH-100 meter, LICHEN Sci-Tech, Co., Ltd., Shanghai, China). Water samples were collected once a week and passed thought a 0.22-μm pore size microfilter (Xinya purification equipment Co., Ltd, Shanghai, China). Then, the dissolved total nitrogen in water (DTN), total ammonia nitrogen (TAN), nitrite, nitrate, carbonate alkalinity, and total suspended solid (TSS), were determined according to the standard methods (APHA, 1995). Additionally, the whole total nitrogen (WTN) in water body of each treatment was also determined every week with water sample without filtration (APHA, 1995). And the total nitrogen contained in biofloc (BTN) was defined as the total nitrogen content contained in biofloc in per liter water and was determined by the difference between WTN and DTN. With an Imhoff cone, 1-liter water sample was settled for 15 min after which the sediment volume was read as the biofloc volume (settleable solids, BFV) (Avnimelech, 2015), and the supernatant was used to measure the turbidity (APHA, 1995).

### 2.4 Growth performance measurements

Thirty PL were selected randomly and individually weighed to the nearest 0.1 mg with an electric balance (AUX220, Shimadzu, Japan) each week. Survival rate, weekly increment of body weight (wiW), specific growth rate (SGR), feed conversion rate (FCR) and productivity were calculated according the following formulates:

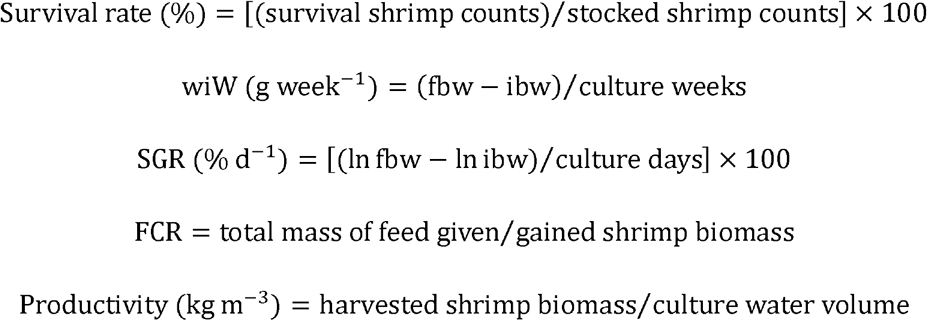

Wherein, fbw and ibw represented the final and initial mean body weight of PL, respectively.

### 2.5 High-throughput sequencing of 16S rRNA gene and data processing

Along the experimental period, water was sampled every week. Briefly, 50 ml water was collected from each tank of a treatment according to the method of Martínez-Córdova et al. (2018), pooled as one sample and filtered through a 0.22-μm pore size membrane. Then, the membrane was stored at −80 °C for extraction of bacterial DNA genome with an E.Z.N.A^TM^ Mag-Bind Soil DNA Kit (OMEGA Bio-Tek, Inc., GA, USA), according to the manufacturers’ instructions.

The V3-V4 region of 16S rRNA gene of bacterial DNA genome was amplified by using the universal primers, 341F: 5’-CCTACGGGNGGCWGCAG-3’ and 805R: 5’-GACTACHVGGGTATCTAATCC-3’, in a 30-μl mixtures containing microbial DNA (10 ng/μl) 2 μl, forward primer (10 μM) 1 μl, reverse primer (10 μM) 1 μl, and 2X KAPA HiFi Hot Start Ready Mix 15 μl (TaKaRa Bio Inc., Japan), via a procedure with a thermal instrument (Applied Biosystems 9700, USA): 1 cycle of denaturing at 95°C for 3 min, first 5 cycles of denaturing at 95°C for 30 s, annealing at 45°C for 30 s, elongation at 72°C for 30 s, then 20 cycles of denaturing at 95°C for 30 s, annealing at 55°C for 30 s, elongation at 72°C for 30 s and a final extension at 72°C for 5 min. The PCR product was purified and recovered with MagicPure Size Selection DNA Beads (TransGen Biotech Co., Ltd., Beijing, China). And then, the purified PCR product was quantified and normalized with a Qubit ssDNA Assay Kit (Life Technologies, USA), to construct the 16S gene library for high-throughput sequencing on the Miseq platform according to the standard protocol (Illumina, USA) by Sangon Biotech (Shanghai) Co., Ltd. The raw data produced from high-throughput sequencing has been deposited at NCBI with accession number of PRJNA646765.

Data processing for the high-throughput sequencing data was carried out under the QIIME 2 (Quantitative Insights Into Microbial Ecology, Version 2019.10) framework (Bolyen et al., 2019). In brief, ambiguous nucleotides, adapter sequences and primers contained in reads, and short reads with length less than 30 bp were removed with the cutadapt plugin (Martin, 2011). After that, bases in the two ends of reads with quality score lower than 25 were trimmed. Thereafter, chimeras were filtered, and pair-ended reads were joined, dereplicated, to obtain high-quality reads which were clustered to operational taxonomic units (OTUs) with an identity of 0.97 by using the Vsearch tool (Rognes et al., 2016). Thereafter, the counts of OTUs were normalized by 16S rRNA gene copy number based on rrnDB database (version 5.6) with the QIIME 2 plugin of q2-gcn-norm.

### 2.6 Bacterial diversity analysis

Bacterial community diversity was analyzed with Plugin diversity of QIIME 2 basing on the normalized counts of OTUs. Metrics regard to alpha diversity, such as Shannon index, richness indexes of Margalef, dominance indexes of Berger-Parker and Simpson, and evenness indexes of Heip and Pielou, were computed with the diversity plugin of QIIME 2. Besides, beta Jaccard index (distance) were also determined with this plugin, and then significantly tested by using PERMANOVA with 999 permutations (Anderson, 2001). Basing on the OTUs composition of samples, each sample was plotted into a PCA diagram with STAMP (Parks et al., 2014).

### 2.7 Bacterial composition analysis

OTUs were annotated by referring to the GreenGene database 13.8, collapsed at phylum, class, order, family and genus levels, respectively, with a QIIME 2 Plugin of taxa. At each level, relative frequencies of species were calculated with the normalized counts of OTUs. Differential analysis for phylum proportion between both treatments was conducted with STAMP (Parks et al., 2014) and LEfSe (Segata et al., 2011).

### 2.8 Statistical analyses

Data was statistically analyzed with the SPSS platform for windows (version 22.0, IBM Co., NY, USA). Student’s T Test were executed for comparison of growth parameters between both treatment, as soon as normality distribution and homogeneity of data were proved with Shapiro-Wilk’s test and Levene’s test, respectively. Or else, non-parametric Mann-Whitney U test was conducted. Percentage data was submitted to arcsine transformation. Furthermore, during each stage, ANOVA with repeated measurements was used to test the significance of main effects and interactions of substrate and time on water parameters, bacterial diversity indices and bacterial composition data under Mauchly’s test of Sphericity assuming condition, or else, the *P* value was corrected by using Greenhouse-Geisser test. Differences were considered significant at *P* < 0.05.

## 3 RESULTS

### 3.1 Water quality

The levels of dissolved oxygen and temperature in the present study were above 5.0 mg L^−1^ and 26.0 °C, respectively (Table 1). The effects of time on both parameters were significant (*P* < 0.05, Table 1). Whereas, substrate showed no significant effect on the levels of dissolved oxygen and temperature (*P* > 0.05, Table 1).

**Table 1.** Water parameters in the biofloc systems with (sB) and without (nB) addition of elastic solid packing filler (nylon) as substrate, during the 28-days experiment culturing *Penaeus vannamei* at a salinity of 5.0‰

| parameters | Treatments |  | *P*-value |  |  |
| --- | --- | --- | --- | --- | --- |
|  | nB | sB | Substrate | Time | Interaction |
| TSS (mg L<sup>-1</sup>) | 148.5±31.3 | 491.3±150.2 | 0.009 | 0.116 | 0.594 |
| WTN (mg L<sup>-1</sup>) | 76.8±11.2 | 76.3±9.6 | 0.057 | <0.001 | 0.013 |
| DTN (mg L<sup>-1</sup>) | 13.0±2.6 | 14.7±3.6 | 0.890 | <0.001 | <0.001 |
| BTN (mg L<sup>-1</sup>) | 63.8±9.3 | 61.6±12.5 | 0.052 | <0.001 | 0.085 |
| TAN (mg L<sup>-1</sup>) | 0.55±0.25 | 0.43±0.06 | 0.010 | 0.227 | 0.002 |
| Nitrite (mg L<sup>-1</sup>) | 0.37±0.28 | 0.10±0.03 | 0.352 | 0.348 | 0.342 |
| Nitrate (mg L<sup>-1</sup>) | 0.51±0.14 | 0.56±0.08 | 0.924 | 0.274 | 0.252 |
| Biofloc volume (mL L<sup>-1</sup>) | 8.8±2.3 | 8.3±2.3 | 0.044 | <0.001 | <0.001 |
| Turbidity (NTU) | 111.9±56.3 | 299.5±92.0 | 0.002 | 0.005 | 0.059 |
| CAK (mg L<sup>-1</sup> CaCO<sub>3</sub>) | 373.7±12.4 | 306.5±51.7 | <0.001 | <0.001 | <0.001 |
| Temperature (°C) | 26.9±0.7 | 27.1±0.6 | 0.656 | <0.001 | 0.012 |
| pH | 7.20±0.14 | 7.11±0.03 | 0.001 | 0.001 | <0.001 |
| Dissolved oxygen (mg L<sup>-1</sup>) | 5.85±0.32 | 5.63±0.59 | 0.215 | <0.001 | 0.764 |

Note: BTN, the total nitrogen contained in biofloc; CAK, carbonate alkalinity; DTN, the dissolved total nitrogen in water; NTU, nephelometric turbidity units; TAN, total ammonia nitrogen; TSS, total suspended solids; WTN, the whole total nitrogen in water.

The carbonate alkalinity in sB treatment was maintained at a high level of 306.5±51.7 mg L^−1^ CaCO_3_, although it was lower than that in nB treatment (373.7±12.4 mg L^−1^ CaCO_3_, Table 1). The pH values in nB treatment and sB treatment were 7.20±0.14 and 7.11±0.03, respectively (Table 1). Both parameters were significantly affected by substrate and time, as well as their interactions (*P* < 0.05, Table 1).

The mean concentrations of the three inorganic nitrogen compounds were at low levels in the present study (< 1.0 mg L^−1^, Table 1). Those compounds were not significantly affected by substrate, time and their interaction (*P* > 0.05), with exceptions of the substrate main effect and the interaction of substrate and time on TAN (*P* < 0.05, Table 1).

No significant substrate effect on WTN (whole total nitrogen in waterbody), BTN (total nitrogen contained in biofloc in per liter water) and DTN (dissolved total nitrogen in waterbody) was found (*P* > 0.05, Table 1). Whereas, the effects of time on those parameters were significant, as well as the interactions of substrate and time on WTN and DTN (*P* < 0.05, Table 1).

The biofloc volume (BFV) levels in nB treatment and sB treatment were similar, with a value of 8.8±2.3 and 8.3±2.3 mL L^−1^, respectively (Table 1). However, the total suspended solids (TSS) and turbidity in sB treatment (491.3±150.2 mg L^−1^, and 299.5±92.0 nephelometric turbidity units, NTU) were higher than those in nB treatment (148.5±31.3 mg L^−1^ and 111.9±56.3 NTU, Table 1). Substrate showed significant main effects on those three parameters (*P* < 0.05, Table 1). The effects of time on BFV and turbidity were also significantly, as well as the interaction of substrate and time on BFV (*P* > 0.05, Table 1).

### 3.2 Growth performance

During the 28-days culture experiment, the average body weights of shrimp in sB treatment were higher than those of nB treatment (Fig. 1). At the end, although the final body weight in sB treatment (0.40±0.03 g) was not significantly different from that of B treatment (0.36±0.04 g, *P* = 0.596, Table 2), the survival rate (96.3±3.6%) and productivity (1.54±0.12 kg m^−3^) of the former treatment were significantly higher than those of the latter (81.0±7.1% and 1.14±0.09 kg m^−3^, *P* < 0.05, Table 2). The feed conversion rate (FCR) in sB treatment (0.76±0.06) was significantly lower than that in nB treatment (0.98±0.08, *P* = 0.044, Table 2). There no significant difference was observed between both treatments for the weekly increment of body weight (wiW) and specific growth rate (SGR) of shrimp during the culture experiment (*P* > 0.05, Table 2).

**Fig. 1.**
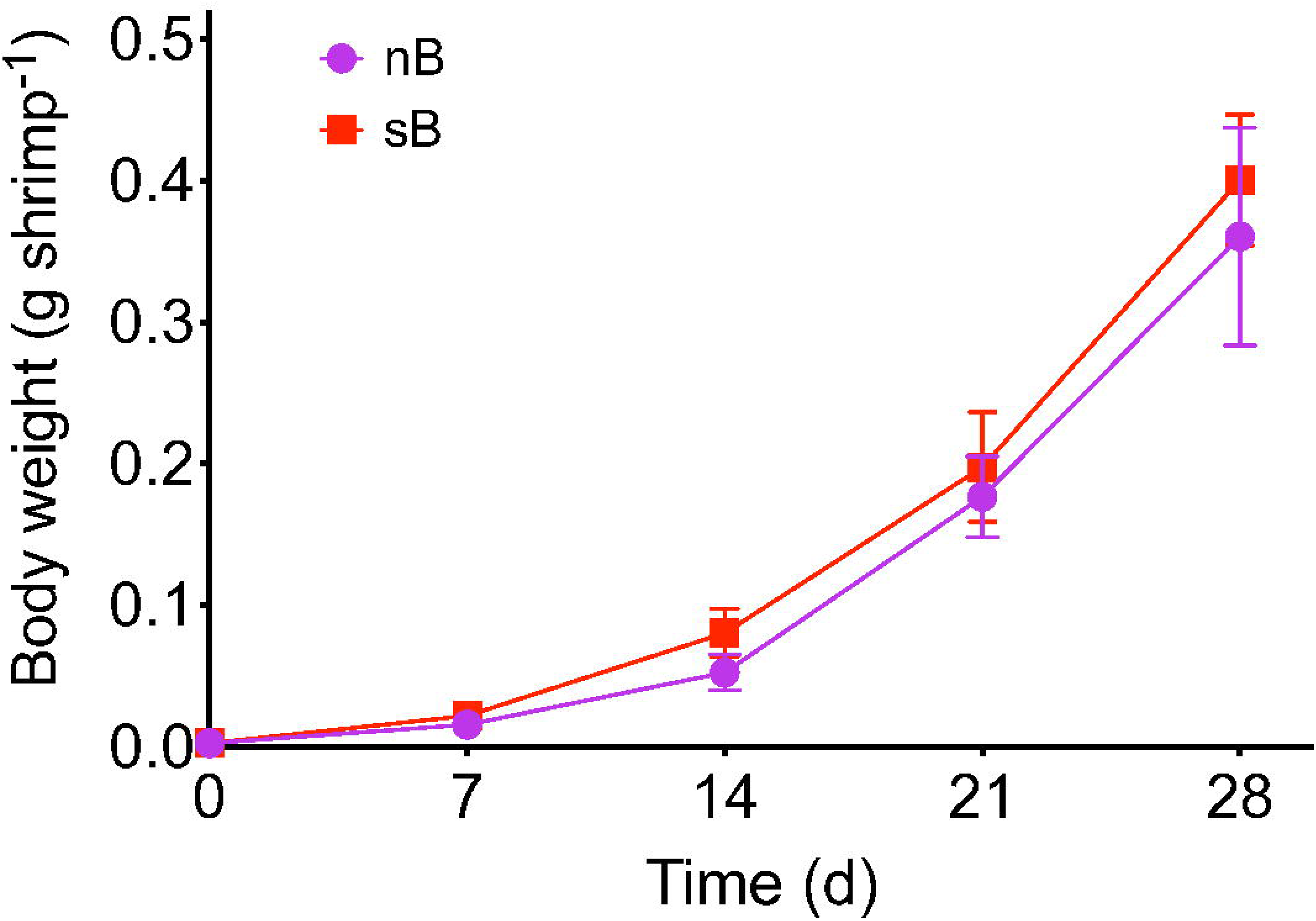
Mean body weight of *Penaeus vannamei* postlarvae cultured in the biofloc systems with (sB) and without (nB) addition of elastic solid packing filler (nylon) as substrate, with time elapsing during the 28-days experiment at a salinity of 5.0‰. Error bar indicates ± standard deviation (SD).

**Table 2.**
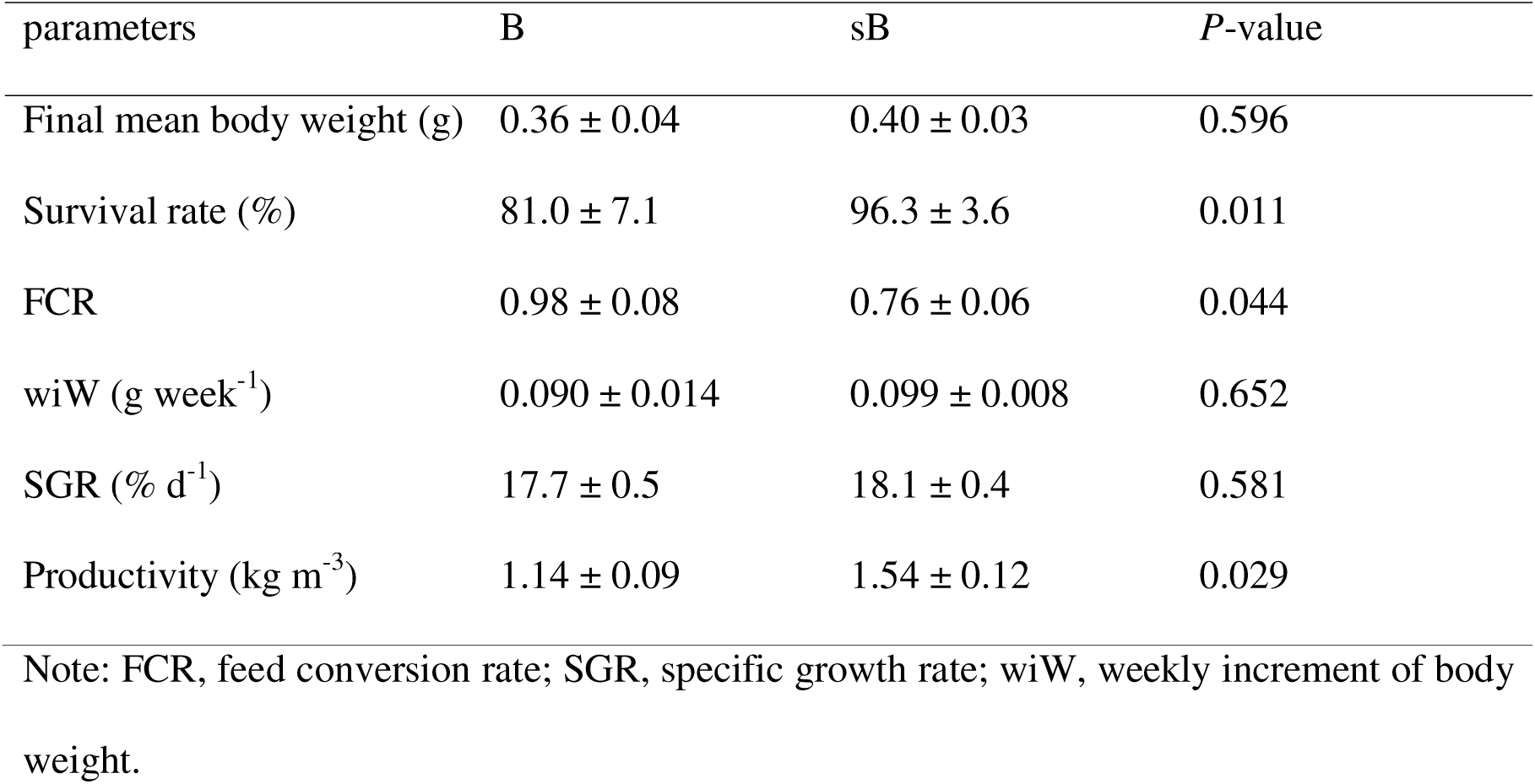
Growth performance of *Penaeus vannamei* postlarvae cultured in the biofloc systems with (sB) and without (nB) addition of elastic solid packing filler (nylon) as substrate, during the 28-days experiment at a salinity of 5.0‰

| parameters | B | sB | *P*-value |
| --- | --- | --- | --- |
| Final mean body weight (g) | 0.36 ± 0.04 | 0.40 ± 0.03 | 0.596 |
| Survival rate (%) | 81.0 ± 7.1 | 96.3 ± 3.6 | 0.011 |
| FCR | 0.98 ± 0.08 | 0.76 ± 0.06 | 0.044 |
| wiW (g week<sup>-1</sup>) | 0.090 ± 0.014 | 0.099 ± 0.008 | 0.652 |
| SGR (% d<sup>-1</sup>) | 17.7 ± 0.5 | 18.1 ± 0.4 | 0.581 |
| Productivity (kg m<sup>-3</sup>) | 1.14 ± 0.09 | 1.54 ± 0.12 | 0.029 |

Note: FCR, feed conversion rate; SGR, specific growth rate; wiW, weekly increment of body weight.

### 3.3 Bacterial diversity

The Shannon index for bacterial community in sB treatment (6.77±0.18) was higher than that in nB treatment (6.14±0.07, Table 3). Substrate, time and their interaction showed significant effects on this index (*P* < 0.05, Table 3). The Shannon index in sB treatment obtained the highest value at 7 d and then slightly decreased, but that in nB treatment peaked at 21 d (Fig. 2 a).

**Fig. 2.**
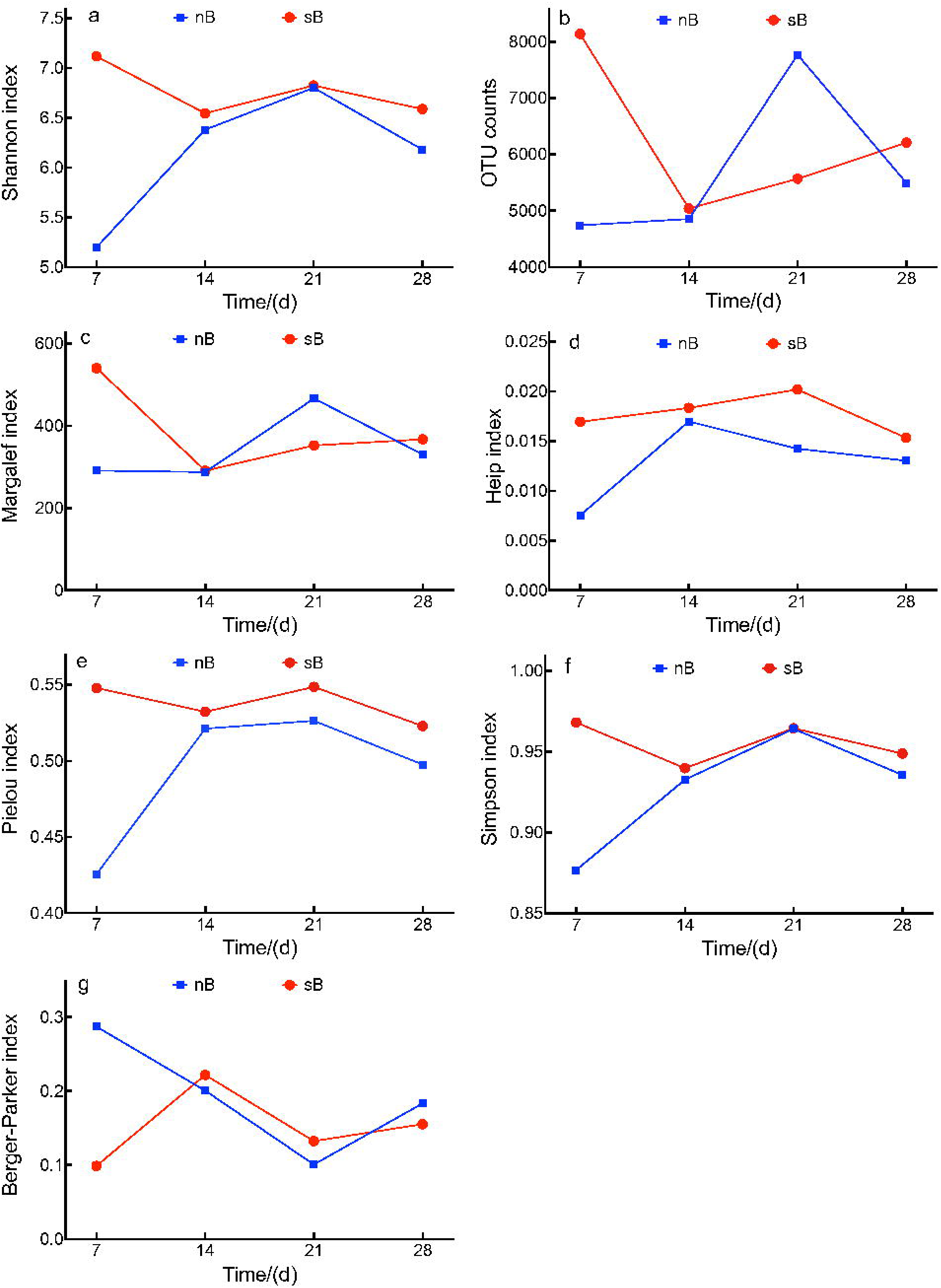
Alpha diversity indexes of bacterial communities in the biofloc systems with (sB) and without (nB) addition of elastic solid packing filler (nylon) as substrate, with time elapsing during the 28-days experiment culturing *Penaeus vannamei* at a salinity of 5.0‰.

**Table 3.** Bacterial alpha diversity indexes in the biofloc systems with (sB) and without (nB) addition of elastic solid packing filler (nylon) as substrate, during the 28-days experiment culturing *Penaeus vannamei* at a salinity of 5.0‰

| Indexes | Treatments |  | P-value |  |  |
| --- | --- | --- | --- | --- | --- |
|  | B | sB | Substrate | Time | Interaction |
| Shannon | 6.14±0.18 | 6.77±0.07 | 0.002 | <0.001 | <0.001 |
| OTU counts | 5713.5±368.2 | 6238.3±353.8 | 0.427 | 0.024 | 0.572 |
| Margalef | 343.9±22.0 | 387.7±28.0 | 0.367 | 0.007 | 0.471 |
| Heip | 0.013±0.001 | 0.018±0.001 | 0.026 | 0.012 | 0.544 |
| Pielou | 0.493±0.012 | 0.538±0.003 | 0.024 | <0.001 | 0.016 |
| B-P | 0.193±0.020 | 0.152±0.014 | 0.098 | 0.002 | 0.676 |
| Simpson | 0.927±0.010 | 0.955±0.003 | 0.043 | <0.001 | 0.127 |

OTU, operational taxonomic unit; B-P, Berger-Parker index.

Higher OTU counts (6238.3±353.8) and Margalef index (387.7±28.0) was observed in sB treatment, when compared to that in nB treatment (5713.5±368.2 and 343.9±22.0, Table 3). Both indexes significantly affected by time (*P* < 0.05), but not by substrate (*P* > 0.05, Table 3). Those two indexes showed similar changing trends with the Shannon index in each treatment (Fig. 2 b and c).

The evenness indexes, Heip index and Pielou index, in sB treatment (0.018±0.001 and 0.538±0.003) and nB treatment (0.013±0.001 and 0.493±0.01, Table 3) peaked at 14 d and 21 d, respectively (Fig. 2 d and e). The main effects and interactions of substrate and time on both indexes were significant (*P* < 0.05), with exception of the interaction on Heip index (*P* = 0.544, Table 3).

The Simpson index in sB treatment (0.955±0.003) was higher than that in nB treatment (0.927±0.010, Table 3). This index in sB treatment displayed a peak value at 14 d, but that in nB treatment showed a decreasing trend during experiment (Fig. 2 f). Substrate and time significantly affected this index (*P* < 0.05, Table 3). Whereas, the Berger-Parker index in sB treatment (0.152±0.014) was lower than that in nB treatment (0.193±0.020, Table 3). Substrate was not significantly affected this index (*P* > 0.05, Table 3). The change trend of Berger-Parker index in the present study was contrary to that of the Simpson index in each treatment (Fig. 2 g).

The bacterial community distances within nB treatment and sB treatment were 0.76±0.36 and 0.77±0.37, respectively (Table 4). PCA analysis also showed that water samples collected from the same treatment were closer than those from the other treatment, basing on the OTUs composition (Fig. 3). In addition, the bacterial beta diversities between both treatments were significantly different with a Jaccard distance of 0.94±0.01 (*P* = 0.001, PERMANOVA, Table 4).

**Fig. 3.**
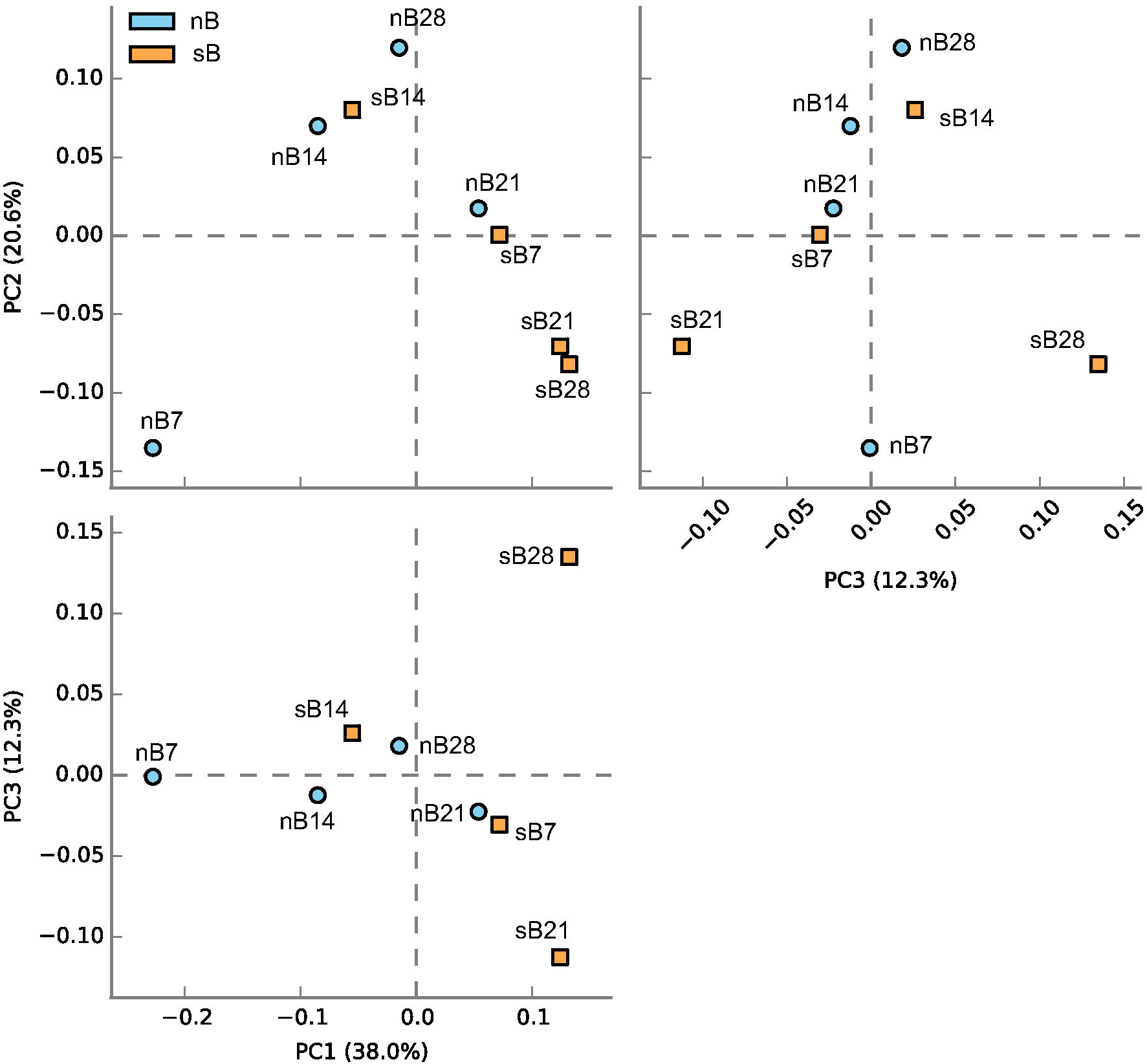
PCA diagram plotting samples basing on their OTU composition profiles. Samples were collected from the biofloc systems with (sB) and without (nB) addition of elastic solid packing filler (nylon) as substrate at time points of 7, 14, 21 and 28 d during experiment culturing *Penaeus vannamei* at a salinity of 5.0‰.

**Table 4.** Bacterial community distance between the biofloc systems with (sB) and without (nB) addition of elastic solid packing filler (nylon) as substrate, during the 28-days experiment culturing *Penaeus vannamei* at a salinity of 5.0‰

| Group 1 | Group 2 | Jaccard distance | Permutations | pseudo-*F* | *P*-value |
| --- | --- | --- | --- | --- | --- |
| B | B | $0.761 \pm 0.361$ | - | - | - |
| sB | sB | $0.768 \pm 0.365$ | - | - | - |
| B | sB | $0.941 \pm 0.007$ | 999 | 3.96 | 0.001 |

### 3.4 Bacterial composition profile

Thirty phyla were assigned for the OTUs in the present study. The most dominant phyla in both treatments were Actinobacteria (4.0-22.7%), Bacteroidetes (10.4-33.5%), Firmicutes (0.2-11.2%), Planctomycetes (4.0-14.9%) and Proteobacteria (29.4-59.0%) (Fig. 4 a). Besides, Chloroflexi and Verrucomicrobia were one of the dominant phyla in nB treatment (0.1-11.4%) and sB treatment (0.3-16.2%), respectively (Fig. 4a). Among those seven phyla mentioned above, the proportions of Chloroflexi, Planctomycetes and Verrucomicrobia were significantly different between both treatments from a point of view of the whole experimental period (*P* < 0.05, Fig. 4 b). The main effects and interactions of substrate and time on proportions of those seven phyla were significant (*P* < 0.05), except the substrate effects on Firmicutes and Proteobacteria, and the interaction of substrate and time on Planctomycetes (*P* > 0.05, Table 5).

**Fig. 4.**
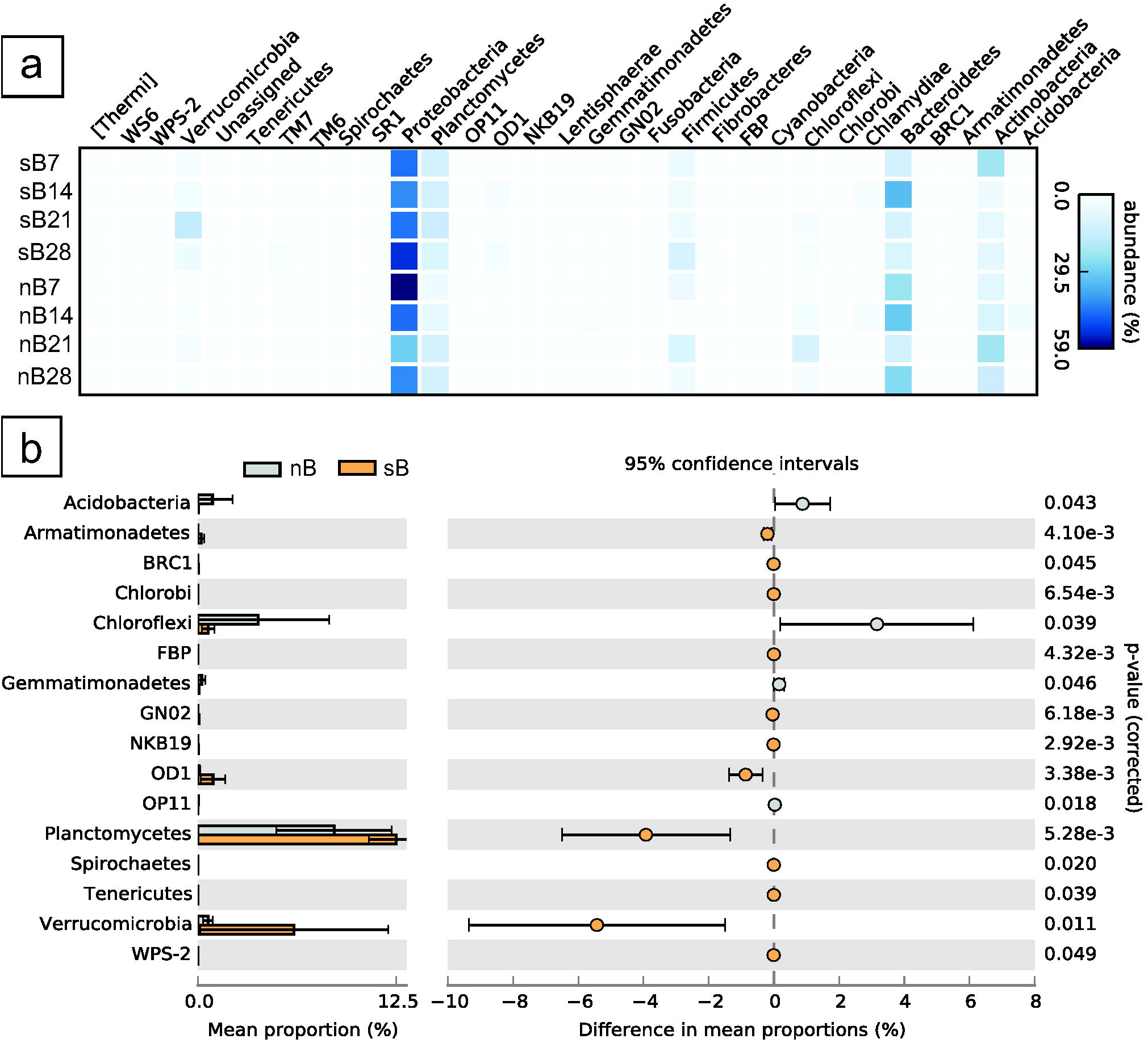
Heatmap displaying bacterial composition profiles at phylum level (a) and differential phyla (b) in the biofloc systems with (sB) and without (nB) addition of elastic solid packing filler (nylon) as substrate, with time elapsing during the 28-days experiment culturing *Penaeus vannamei* at a salinity of 5.0‰.

**Table 5.** Proportions (%) of dominant phyla in the biofloc systems with (sB) and without (nB) addition of elastic solid packing filler (nylon) as substrate, during the 28-days experiment culturing *Penaeus vannamei* at a salinity of 5.0‰

| Phylum | Treatments |  | P-value |  |  |
| --- | --- | --- | --- | --- | --- |
|  | nB | sB | Substrate | Time | Interaction |
| Actinobacteria | 13.67±3.34 | 10.00±4.26 | <0.001 | <0.001 | <0.001 |
| Bacteroidetes | 23.18±4.01 | 16.87±5.57 | <0.001 | <0.001 | <0.001 |
| Chloroflexi | 3.79±2.57 | 0.63±0.22 | <0.001 | 0.001 | 0.002 |
| Firmicutes | 4.53±2.09 | 5.92±1.79 | 0.087 | 0.001 | 0.010 |
| Planctomycetes | 8.57±2.10 | 12.49±1.00 | 0.006 | 0.035 | 0.486 |
| Proteobacteria | 43.82±6.10 | 45.90±2.46 | 0.405 | <0.001 | 0.038 |
| Verrucomicrobia | 0.63±0.18 | 6.06±3.42 | <0.001 | <0.001 | <0.001 |

The bacterial composition profiles on class, order, family and genus levels were showed in Fig. 1S–Fig. 4S, respectively.

The LEfSe analysis showed that an unassigned genus and the phylum Verrucomicrobia were the most significant biomarkers for nB and sB, respectively (Fig. 5).

**Fig. 5.**
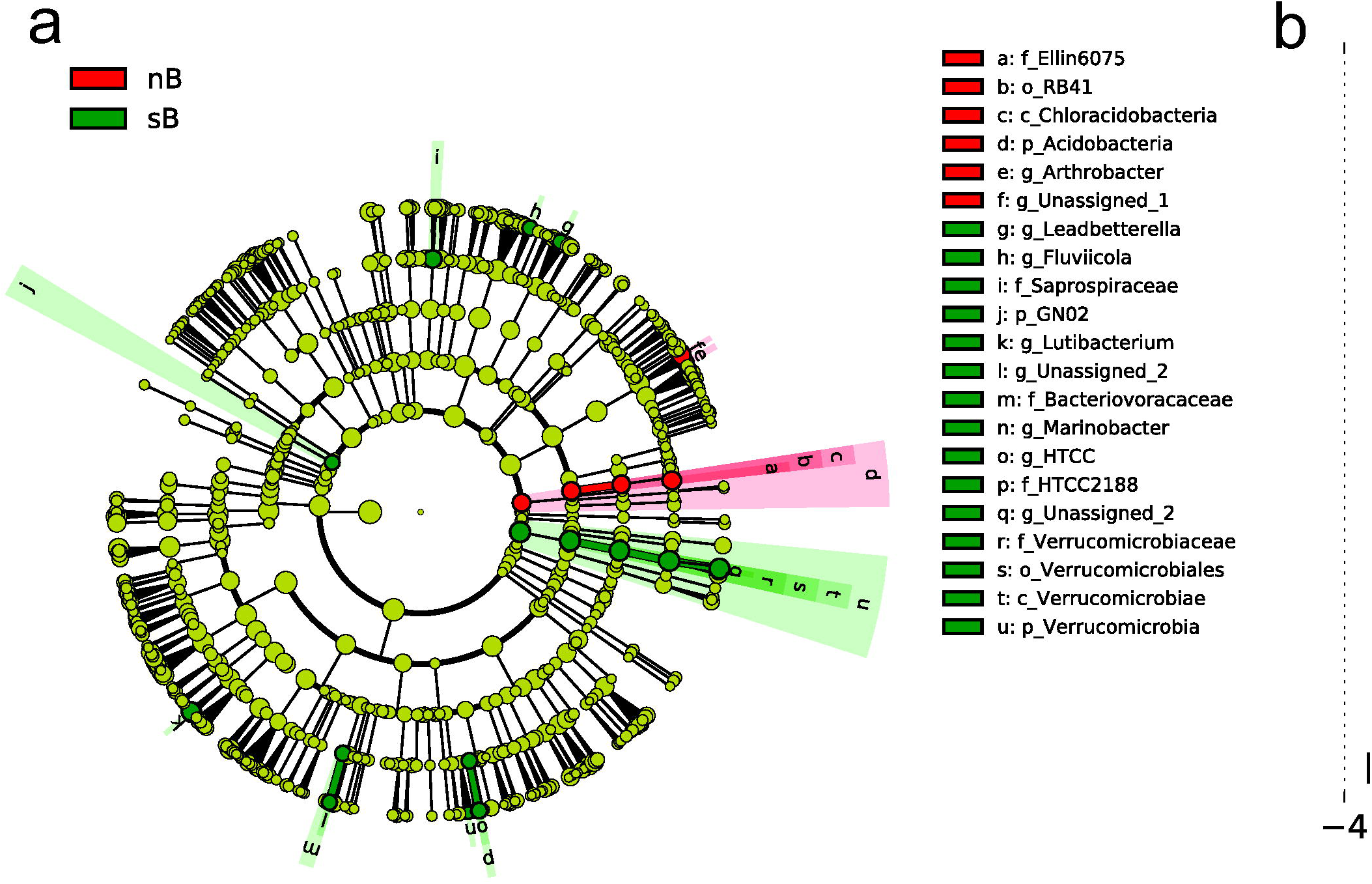
Cladogram (a) and Linear discriminant analysis (LDA) scores of LEfSe (linear discriminant analysis effect size) analysis showing biomarkers in the biofloc systems with (sB) and without (nB) addition of elastic solid packing filler (nylon) as substrate, with time elapsing during the 28-days experiment culturing *Penaeus vannamei* at a salinity of 5.0‰.

## 4 DISCUSSION

### 4.1 Effects of substrate on water quality

All the water parameters determined in the present study were within the acceptable range for culturing *P. vannamei* (Furtado et al., 2014; Hargreaves, 2013; Schveitzer et al., 2013a; Van Wyk et al., 1999; Xu et al., 2012; Zhang et al., 2017). Addition of substrate showed significantly effect on TSS in the present study (*P* < 0.05), with higher level in sB treatment than that in nB treatment. Consistently, Schveitzer et al. (2013b) also observed significant effect of substrate on TSS with higher levels in the biofloc systems with substrate. Whereas, in another previous study, the TSS levels in the biofloc systems with substrate were lower than that in the system without substrate (Rezende et al., 2018). It is proposed here that substrates with different shape have different effects on TSS. For example, nylon or polyester fiber net with a small mesh size (1-2 mm) has the ability to filter the water and capture biofloc, thereby reducing the amount of solids in the rearing water, leading to the lower TSS value (Rezende et al., 2018). On the contrary, the substrate with a larger mesh size, such as 2.5 cm (Tierney et al., 2020), would filter both water and biofloc. Similarly, in the present study, elastic solid packing filler (nylon) was used as substrate which is a long open space composed with nylon yarns radially fixed to a central rope. Thus, the flocs would be not retained within this open space of substrate, especially under the rough aeration condition of the biofloc system, leading to an increasing of TSS level in water body.

In the present study, it was found that the turbidity levels in both treatments were increased continuously. This might be contributed to the removal operation for biofloc during 14-28 d of the experiment in the present study. The turbidity in the present study was determined on water sample without big-size biofloc which settled by using an Imhoff cone for 15 min, indicating that this parameter was correlated to the content of small-size biofloc. Previous studies have showed that the bacterial groups are different between small- and big-size bioflocs (Chen et al., 2019; Huang et al., 2020). Therefore, the removal operation in this study might make settleable solids (big-size biofloc) as well as bacterial groups attached be removed from the water column, which would improve the growth of bacteria adhering on small-size biofloc retained in the water body, by reducing the competition from bacteria relating with big-size biofloc, and in turn, prompting formation of small-size suspended biofloc and increasing the turbidity. Substrate showed significant effect on turbidity in the present study (*P* = 0.002). It was thought that the addition of artificial substrates may influence the water circulation in the tanks, leading to smaller turbulence in the water, and in turn to facilitating sedimentation or particle aggregation and formation of large flocs, which reduces the suspended solids and thus the turbidity in the water column (Ferreira et al., 2016; Fleckenstein et al., 2020). However, in the present study, the turbidity level in sB treatment was found to be higher than that in nB treatment. It was speculated that substrate might play a positive role in the process of turbidity increasing caused by the removal operation for big-size biofloc discussed above.

Substrate also showed significant effect on carbonate alkalinity in the current study (*P* < 0.001). And the carbonate alkalinity in sB treatment was lower than that in nB treatment in the current study, indicating that more alkalinity was consumed in biofloc system with substrate. This might be contributed to the different bacterial communities which led to different bacterial activities consuming HCO_3_^−^ (Ebeling et al., 2006). Corroboratively, more diverse bacterial community in sB treatment (Shannon index 6.77±0.18) than that in nB treatment (Shannon index 6.14±0.07), and different bacterial communities between both treatments (Jaccard distance 0.94±0.01, *P* < 0.001) were observed in the present study. Nevertheless, the inferring function profile of bacterial community should be investigated to better understand the difference on HCO_3_^−^ consumption between both treatments next.

The results of ANOVA with repeated measurements showed that addition of substrate showed significantly effects on TAN, biofloc volume (BFV), and pH in the present study (*P* < 0.05). Meanwhile, the interactions of substrate and time on most water parameters, such as whole total nitrogen in waterbody (WTN), total dissolved nitrogen (DTN), TAN, BFV, carbonate alkalinity and pH, were significant in the present study, indicating that the changing styles of these parameters along with time elapsing were different between the biofloc systems with and without addition of substrate. Rezende et al. (2018) also observed that some water parameters were significantly affected by the interactions of the rearing day and treatment (substrate).

### 4.2 Effects of substrate on shrimp growth

The body weight of shrimp in treatment with substrate would be significantly prompt, when compared to that without substrate (Olier et al., 2020; Schveitzer et al., 2013b), via forming periphytons on substrate as supplementary natural food which could be fed by shrimp for 24 h a day (Anand et al., 2019; Asaduzzaman et al., 2008; Kumar et al., 2017). In the present study, the final body weight, weekly increment of body weight (wiW) and specific growth rate (SGR) of shrimp in sB treatment were elevated by 10.0-11.1%, compared to those in nB treatment, which were similar with the results (8.4-11.0%) observed by Tierney et al. (2020). Previous studies showed that the raw material (Mani et al., 2021; Peiro-Alcantar et al., 2019; Rezende et al., 2018) and shape (Fleckenstein et al., 2020) of substrates could affect its elevating efficiency on growth performance of shrimp. For example, periphyton nutritional values were shown to be dependent on substrate type (Tinh et al., 2021). And natural or vegetable substrates was found to be more beneficial to growth of shrimp than artificial substrates (Mani et al., 2021). It is also thought that the substrate composed of individual sheets with a large opening or mesh size and a low overall surface area would show less improving effect on shrimp production metrics. For instance, Ferreira et al. (2016) found that the addition of substrate with an amount equivalent 400% of the lateral area of the tanks significantly elevated the body weight of shrimp, but that with 200% did not. Similarly, Fleckenstein et al. (2020) and Tierney et al. (2020) also observed that addition of substrate only increased the total in-tank surface area by 13.4% and 21%, respectively, did not lead to significant effect on any shrimp production metrics. In this study, the substrate used was a long open space composed with nylon yarns radially fixed to a central rope, covering only 52.4% of the tank internal surface, which might be contributed to the non-significant benefit for the final body weight, wiW and SGR of shrimp in sB treatment, compared to those in nB treatment (*P* > 0.05).

Nevertheless, in the present study, the survival rate and productivity in sB treatment significantly increased when compared with those in nB treatment (*P* < 0.05). Substrate could act as a refuge to add space and reduce the cannibalism behavior, leading to more survival. And this increased space by addition of substrate reduced the relative stocking density, which is beneficial for shrimp growth. An inverse relationship between stocking density and growth performance of shrimp was found in previous studies (Esparza-Leal et al., 2015; Esparza-Leal et al., 2020; Liu et al., 2017; Panigrahi et al., 2019; Peixoto et al., 2018). Moreover, the feed conversion rate (FCR) in sB treatment significantly decreased, compared with that in nB treatment (*P* < 0.05). Bioflocs which mostly grew on tank wall or substrate as periphyton could be grazed by shrimp as natural supplementary food (Tinh et al., 2021), reducing input of artificial feeds, and consequently decreasing the FCR (Avnimelech, 2015). Undoubtedly, substrate addition facilitates this feeding behavior of shrimp by supplying a niche for preying on biofloc (Anand et al., 2019; Asaduzzaman et al., 2008).

### 4.3 Effects of substrate on diversity of bacterial diversity

Generally, highly diverse bacterial community including 18-40 phyla were observed in waterbody of biofloc systems raising *P. vannamei* (Baliga et al., 2021; Ferreira et al., 2021; Huerta-Rabago et al., 2019; Vargas-Albores et al., 2019), with a Shannon index as high as 5.31-7.23 (Addo et al., 2021; Chen et al., 2019; Gao et al., 2019; Huang et al., 2021; Jiang et al., 2020; Kim et al., 2021; Pilotto et al., 2018). Similarly, in the present study, 30 phyla were observed in both treatments with a Shannon index of 6.14-6.77. In the biofloc system, the environment rich in organic matter could form several trophic levels which facilitate the growth of a number of bacterial phyla (Fleckenstein et al., 2020), resulting in a high bacterial diversity.

Between both treatments in the present study, the beta diversity Jaccard index was significantly different, indicating different bacterial communities in both treatments. Corroboratively, from a point of view of the whole experimental period, higher Shannon index, OTU counts, Margalef index, Heip index, Pielou index and Simpson index, but lower Berger-Parker index of bacterial community in sB treatment were observed when compared to those in nB treatment, indicating higher bacterial diversity, richness and evenness, but lower dominance of predominant species (Magurran, 2004), in the biofloc system with addition of substrate (sB treatment).

The results of ANOVA with repeated measurements revealed that substrate showed significant effects on some alpha diversity indexes, such as Shannon index, Heip index, Pielou index and Simpson index (*P* < 0.05). Meanwhile, the interactions of substrate and time on Shannon index and Pielou index were also significant (*P* < 0.05), indicating that the changing styles of those indexes over time were different between treatments in the present study. Consistently, the bacterial community in sB treatment obtained the most diverse situation at 7 d with the highest Shannon index, but in nB treatment, the Shannon index increased gradually and peaked at 21 d, indicating that substrate addition accelerated the formation of high-diverse bacterial community in the biofloc system.

All alpha diversity indexes were significantly affected by time (*P* < 0.05), indicating a time-dependent succession pattern of bacterial diversity in biofloc systems regardless of addition of substrate, which was also observed in other studies (Jiang et al., 2020; Kim et al., 2021; Wang et al., 2020; Xu et al., 2019).

### 4.4 Effects of substrate on bacterial composition

In general, Proteobacteria and Bacteroidetes were found to be the most dominant phyla in marine biofloc systems rearing *P. vannamei* (Ferreira et al., 2021; Huerta-Rabago et al., 2019; Jiang et al., 2020; Pilotto et al., 2018; Martínez-Córdova et al., 2018; Vargas-Albores et al., 2019), as well as in biofloc systems with low salinities (Addo et al., 2021; Huang et al., 2021). Similarly, in the present study, those two phyla together accounted for 41.33-82.44% and 55.56-74.69% in nB treatment and sB treatment, respectively. Proteobacteria have a diverse phenotypic and phylogenetic lineage and widely dispersed in the environment rich in organic compounds (Kersters et al., 2006; Kirchman, 2002; Rud et al., 2017), such as biofloc systems. Bacteroidetes serve as a degrader of polymeric organic matter and are found in a range of habitats (Thomas et al., 2011; Woebken et al., 2007). Additionally, Actinobacteria, Chloroflexi, Firmicutes, Planctomycetes and Verrucomicrobia were important phyla in both treatments of the present study, which were also commonly detected in biofloc systems stocking white-leg shrimp (Deng et al., 2018; Ferreira et al., 2021; Huang et al., 2021; Huerta-Rabago et al., 2019; Jiang et al., 2020; Luis-Villaseñor et al., 2016; Martínez-Córdova et al., 2018; Martins et al., 2020; Schveitzer et al., 2020).

Substrate showed significantly main effects on the proportions of most of the seven predominant phyla mentioned above in this current study. It is considered that substrate could supply attaching material for species which grow adhesively, and tend to adsorb the bacteria with particle-attaching or biopolymer-degrading capability (Huang et al., 2020). For example, the filamentous Chloroflexi grow on other microorganisms and substrate (Kragelund et al., 2007; Miura et al., 2007); and Bacteroidetes prefer for polymers and are frequently found attaching to surfaces, colonizing on particles (Deng et al., 2019; Fernández-Gómez et al., 2013; Kirchman, 2002; Morgan-Sagastume et al., 2008). Although Planctomycetes grow as single free-swimming cells, they also appear as single cells within flocs (Morgan-Sagastume et al., 2008), and are usually found to adhere to the surfaces of invertebrates (Fuerst et al., 1997), detrital aggregates (Crump et al., 1999; Delong et al., 1993), and flocs (Huang et al., 2020). Actinobacteria are chemoorganotrophic bacteria (saprophytes) with branched filaments (Madigan et al., 2015) and have been described as small free-living cells (Ghai et al., 2013), leading to a higher proportion in free-living fraction than that in particle-attached fraction (Bachmann et al., 2018; Huang et al., 2020; Mestre et al., 2017). Therefore, the free- or attaching-living characteristic of those phyla might contribute to the variations of their abundance between both treatments in the present study.

Although Firmicutes are microorganisms to grow as single free-swimming cells making them more likely to remain in suspension (Morgan-Sagastume et al., 2008). And Proteobacteria survive as free-living to live suspended in the water column, although some species might attach to particles, surfaces, or cells (Fernández-Gómez et al., 2013; Morgan-Sagastume et al., 2008; Rud et al., 2017; Schveitzer et al., 2020), substrate did not show significant effect on those two phyla in this present study. And the reason should be investigated next.

The dominant phyla observed in the present study have variable functions. For instance, Proteobacteria play important roles in the process of nutrient cycling and the mineralization of organic compounds (Kersters et al., 2006; Kirchman, 2002; Rud et al., 2017); Bacteroidetes produce extracellular enzymes with degradative capabilities to serve as degraders of polymeric organic matter (Kirchman, 2002; Thomas et al., 2011; Woebken et al., 2007); Chloroflexi reductively dehalogenate organic chlorinated compounds to obtain energy (Gupta, 2013); Actinobacteria are important in the nutrient and biomaterial recycling (Goodfellow and Williams, 1983), responsible for degrading nucleic acids and proteins (Madigan et al., 2015), and for the production of many hydrolytic enzymes (chitinase, chitinolytic, fatty acid, cellulolytic, nitrile hydrolyzing, amylolytic, proteolytic, and lipolytic) (Das et al., 2008); Planctomycetes have function to degrade the biopolymers of adsorbate (Woebken et al., 2007). Therefore, the different bacterial community profiles between both treatments in the current study might contribute to different activities of nitrogen and carbon metabolism in their water body. However, the inferring functions of bacterial community should be induced next to directly estimate the effects of addition of substrate on the bacterial activities.

The interactions of substrate and time on proportions of most of those phyla (except Planctomycetes) also significant in the present study, indicating different development styles of bacterial composition profile between both treatments. Furthermore, all the seven dominant phyla in the present study were significantly affected by time (*P* < 0.05), indicating a time-dependent dynamics of bacterial composition in biofloc systems, similar with that observed in other studies (Baliga et al., 2021; Gao et al., 2019; Jiang et al., 2020; Martínez-Córdova et al., 2018; Xu et al., 2019).

## 5 CONCLUSION

In the present study, addition of elastic solid packing filler (nylon, with a diameter of 15 cm) as substrate that coated 52.4% of the tank internal surface in the biofloc system nursing *P. vannamei* PL with a salinity of 5‰ showed positive effects on shrimp growth performance. Furthermore, substrate also showed significant main effects or interactions with time on most water parameters as well as bacterial diversity and composition.

## Acknowledgements

This work is financially supported by the scientific research program of the Education Department of Hunan province (18B394), the open projects of Hunan Provincial Key Laboratory for Molecular Immunity Technology of Aquatic Animal Diseases (2021KJ001), Changde Research Center for Agricultural Biomacromolecule (2020AB08).

## Notes

### Competing Interest Statement

The authors have declared no competing interest.

### Summary of Updates

There are a error in the reference.

